# Testing a non-destructive assay to track *Plasmodium* sporozoites in mosquitoes over time

**DOI:** 10.1101/2023.08.22.554268

**Authors:** Catherine E. Oke, Sarah E. Reece, Petra Schneider

**Affiliations:** Institute of Ecology and Evolution; Institute of Immunology and Infection Research, School of Biological Sciences, University of Edinburgh, Edinburgh, UK

**Keywords:** extrinsic incubation period, *Anopheles stephensi*, *Plasmodium berghei*, *Plasmodium chabaudi*, malaria transmission

## Abstract

**Background:** The extrinsic incubation period (EIP), defined as the time it takes for malaria parasites in a mosquito to become infectious to a vertebrate host, is one of the most influential parameters for malaria transmission but remains poorly understood. The EIP is usually estimated by quantifying salivary gland sporozoites in subsets of mosquitoes, which requires terminal sampling. However, assays that allow repeated sampling of individual mosquitoes over time could provide better resolution of the EIP.

**Methods:** We tested a non-destructive assay to quantify sporozoites of two rodent malaria species, *Plasmodium chabaudi* and *Plasmodium berghei*, expelled throughout 24hr windows, from sugar-feeding substrates using quantitative PCR.

**Results:** The assay can quantify sporozoites from sugar-feeding substrates, but the prevalence of parasite positive substrates is low. Multiple methods to increase the detection of expelled parasites (running additional technical replicates; using groups rather than individual mosquitoes) did not increase the detection rate, suggesting that expulsion of sporozoites is variable and infrequent.

**Conclusions:** We reveal successful detection of expelled sporozoites from sugar-feeding substrates. However, investigations of the biological causes underlying the low detection rate of sporozoites (e.g. mosquito feeding behaviour, frequency of sporozoite expulsion, or sporozoite clumping) are needed to maximise the utility of using non-destructive assays to quantify sporozoite dynamics. Increasing detection rates will facilitate the detailed investigation on infection dynamics within mosquitoes, which is necessary to explain *Plasmodium’s* highly variable EIP and improve understanding of malaria transmission dynamics.

## Background

Malaria, caused by *Plasmodium* parasites, [1] is transmitted between vertebrate hosts by *Anopheline* mosquito vectors. Within the vector, parasites must mate, reproduce, traverse the midgut wall, replicate extensively and then migrate to the salivary glands. Only after all these processes (defined as sporogony) are completed, can parasites infect a new vertebrate host. The time it takes for parasites to complete their development in the vector (the extrinsic incubation period, EIP [2]) is usually reported to be 10-20 days [3,4]. This is surprisingly long given that only a very small proportion of mosquitoes live longer than three weeks in the field [5–7].

Small changes in the EIP can have a large effect on the number of mosquitoes living long enough to become infectious, making it a crucial parameter for transmission potential (i.e. R_0_) [3]. Although the historical assumption that the EIP only depends on temperature [8–10] has been overturned [3], understanding of other sources of variation in EIP remains limited. Variation in the EIP is associated with environmental factors, such as temperature and availability of the vector’s resources, along with intrinsic differences between *Plasmodium* species. For example *Plasmodium mexicanum*, vectored by the short-lived sand fly, has a shorter EIP [2], whereas *P. berghei* has a longer EIP, partly due to adaptation to the lower temperature of their vector’s habitat [11]. In comparison, *P. chabaudi* and *P. falciparum* have similar development times, with the latter speeding up when mosquitoes receive an additional blood meal [12]. Furthermore, longer EIPs have been observed in *P. falciparum*-infected mosquitoes with lower salivary gland burdens [13]. Why malaria parasites cannot develop faster is a longstanding mystery and highlights the need to investigate whether constraints (such as the dynamics of resource availability within mosquitoes) and/or benefits to the parasite (such as transmission correlating positively with sporozoite number) shape the EIP [6].

Explaining the EIP is challenging because it is most commonly approximated as the time at which sporozoites are first visible in the salivary glands [3]. However, sporozoites may require a period of maturation to become infectious; heterogeneous gene expression suggests that not all sporozoites residing in the salivary glands are infectious [14]. Additionally, salivary gland sporozoites may need to exceed a density threshold for onwards transmission to be likely. For example, while transmission probability significantly increases above 10000 sporozoites for *P. yoelii* [15], even though only tens to low hundreds of sporozoites are thought to be expelled during transmission [13,16–18], studies using *P. falciparum* suggest a lower (>1000) threshold [19]. Furthermore, some infected mosquitoes do not expel any sporozoites [13,20,21] further complicating the correlation between salivary gland sporozoites and transmission probability.

Tools for estimating the EIP are also problematic; the EIP is typically estimated from terminal sampling of a subset of mosquitoes from the population at intervals during sporogony. Sporozoites are usually assayed following dissection of the salivary glands for microscopic detection of sporozoites [22], or by molecular assays from (bisected) mosquitoes [13,23]. These methods have several limitations. First, terminal sampling prevents tracking individual mosquitoes over time so EIP is estimated at population-level. While population-level measures such as the median EIP (EIP_50_) are useful for modelling purposes [3], they do not take into account the individual variation important for linking vector-parasite-environment interactions with the EIP and infectiousness [24,25]. Second, processing a subset of mosquitoes every few days is laborious and requires large numbers of infected mosquitoes.

Sporozoites are expelled during sugar feeding [26,27] and expelled sporozoites have a greater chance of being infectious than those in the glands. Thus, using a non-destructive assay that quantifies expelled sporozoites on sugar-feeding substrates allows the infections of individual mosquitoes to be followed over time and can improve resolution of the EIP. Non-destructive sugar-based assays to quantify sporozoites, using PCR or immunoblotting detection of circumsporozoite protein, have been tested for groups or single mosquitoes infected with *P falciparum* [24,26–29], and for groups of *P. berghei*-infected mosquitoes [28,30] with some success. While assays able to detect sporozoites from groups of mosquito are useful for field surveillance of malaria prevalence [26–28], assays sensitive enough to detect sporozoites from individual mosquitoes provide the best resolution of EIP and its determinants. Furthermore, investigating the ecological and evolutionary determinants of the EIP requires model systems in which the full life cycle can be manipulated. Due to their tractability, rodent malarias are ideal, but there is no assay available for individual mosquitoes infected with these *Plasmodium* species. The most commonly used model, *P. berghei*, is useful for proof of principle investigation of EIP-related questions, including onward transmission to a vertebrate host, but *P. chabaudi* provides a unique opportunity to investigate EIP at a similar parasite density and temperature [31] to *P. falciparum*.

Here we test a non-destructive method to detect *P. berghei* and *P. chabaudi* sporozoites from mosquitoes’ sugar-feeding substrates. We compare how well this technique performs for *P. berghei* and *P. chabaudi* which have different optimal temperatures for sporogony and therefore different EIPs. We demonstrate that *Plasmodium* DNA from both species can be detected and quantified from sugar-feeding substrates. However, while the detection rate for sporozoites in mosquito expectorates is similar to other studies [24,28], parasite prevalence is low. We discuss potential explanations for low parasite prevalence in individual mosquito’s expectorate, and suggest further improvements to sugar-feeding assays.

## Methods

The qPCR to quantify *Plasmodium* sporozoites was validated, and used to determine the best sugar-feeding substrate for the assay, the range of sporozoite DNA concentrations that can be recovered from the substrates, as well as optimal storage conditions and sugar concentrations to minimise DNA degradation. Subsequently, the recovery of expelled sporozoites from individual mosquitoes was investigated, as well as methods to increase the detection of expelled parasites.

### Mosquitoes and malaria infections

*Anopheles stephensi* SD500 mosquitoes were reared at 26°C, 70% relative humidity, in a 12L:12D hr light cycle, with *ad libitum* access to 8% fructose solution post-emergence. Transmission to mosquitoes was achieved through blood-feeding mosquitoes on mice (8-10 week old male C57Bl/6) with microscopy-confirmed gametocytes of either *P. berghei* ANKA or *P. chabaudi* genotype ER (following [31,32]). Mosquitoes used for *P. berghei* transmissions were gradually acclimatised to 21°C prior to infectious blood feeds. All mosquitoes were starved for 24 hours before infection and unfed females were removed on day 1 post-infectious blood meal (pIBM).

### DNA extraction

DNA from microscopy-quantified blood stage parasites [33] was extracted from 5µL blood using a semi-automatic Kingfisher Flex Magnetic Particle Processor and MagMAX^TM^-96 DNA Multi-Sample Kit (ThermoFisher Scientific) as per [34], and was frozen at -20°C until use. These blood-stage DNA samples were used to determine qPCR efficiency and the limit of detection (LOD).

DNA was extracted from mosquitoes and feeding substrates following the CTAB-based phenol-chloroform extraction method from Chen *et al* [35] with minor modifications as described in Schneider *et al.* [33]. Extracted DNA was eluted in 30µL (mosquitoes, supplemented feeding substrates) or 16µL (mosquito expectorate substrates) water, and frozen at -20°C until use. Mosquito, but not feeding substrate DNA extracts, were diluted 4-fold to reduce the effect of inhibitors originating from mosquito material on the performance of the PCR. Seven microliters of (diluted) DNA extracts were used in all PCR reactions, and data are presented as genomes/PCR to account for differences in sample processing.

### Quantification of *Plasmodium* by quantitative PCR

Both *P. berghei* and *P. chabaudi* were assayed by a quantitative PCR (qPCR) targeting a region of the 18S rRNA gene that is highly conserved among *Plasmodium* species [36]. Quantification of parasite genomes was determined by comparing threshold cycle (Ct) against a standard curve, generated from DNA extracted from blood stage parasites of either *P. berghei* ANKA or *P. chabaudi* genotype ER (see “DNA extraction”). Negative water controls were included to identify false positives.

### Optimising the assay

The assay was optimised using two reference DNA samples from sporozoite-infected mosquitoes. DNA samples from *Plasmodium-*infected mosquitoes, shown by qPCR to have high sporozoite loads were pooled to create one reference DNA sample for *P. berghei* (2967 genomes/µL) and one for *P. chabaudi* (5099 genomes/µL). These reference samples were used to determine: 1) which type of feeding substrate type returned an optimal DNA yield, and whether 2) DNA could be detected across a range of concentrations, 3) DNA degradation occurred during the collection period, and 4) sugar content impacted DNA yield.

The most suitable feeding substrate was selected by comparing the recovery of parasite DNA from 15mg cotton wool, a 1cm^2^ cotton pad (Boots UK) or a 1 cm^2^ filter paper (Whatman No. 1). Each substrate (n=3 per substrate type) was soaked in 8% fructose, supplemented with 5µL *P. berghei* or *P. chabaudi* reference DNA and stored at 26°C and 70% relative humidity for 24hrs to mimic housing conditions for *P. chabaudi*-infected mosquitoes. Reduced DNA yields are expected at higher temperatures [37], so these conditions provide a conservative estimate of assay performance for *P. berghei*. DNA yield was calculated by comparing qPCR results directly from reference DNA with those from spiked feeding substrates, accounting for any dilutions during sample processing. Subsequent tests were conducted using cotton wool, and all substrates were kept in the same conditions as described above. Second, to confirm that DNA could be consistently detected and quantified across a range of concentrations, cotton wool substrates were supplemented with 5µL of serial dilutions of *P. berghei* (neat 10^0^ to 5×10^-3^ dilution) or *P. chabaudi* (neat 10^0^ to 5×10^-4^ dilution) reference samples, (n=3 per dilution/species), immediately after soaking in 8% fructose (time point 0hrs). Linearity of quantification and the limit of quantification (LOQ), relative to the limit of detection (LOD), was quantified. Third, DNA degradation under conditions mimicking mosquito housing was tested by comparing DNA recovery from cotton wool supplemented with 5µL reference sample (10^0^ to 10^-2^ dilution for each species) either immediately after soaking in 8% fructose (time point 0hrs) or at collection (time point 24hrs) (n=3 per time point/species). Finally, the impact of sugar concentration on DNA yield was tested by soaking cotton wool substrates in distilled water, 1% or 8% fructose and supplemented with 5µL of serial dilutions of *P. berghei* or *P. chabaudi* reference samples (neat 10^0^ to 10^-2^ dilution, n=3 per dilution/species). Parasite DNA recovery was compared between the 3 sugar concentrations.

### Testing the assay on mosquito expectorate samples

To collect expectorate samples, mosquitoes were moved to paper cups, either individually (*P. berghei* n=13; *P. chabaudi* n=10) or in groups (*P. berghei,* 4 mosquitoes/cup, n=5 cups). To increase the likelihood of sugar feeding, mosquitoes were starved for 24 hours prior to being provided with a feeding substrate, which was collected 24hrs later and stored at -20°C until DNA extraction. This 2-day starvation–feeding cycle was repeated twice during days 22-25 pIBM for *P. berghei*, and days 12-15 pIBM for *P. chabaudi* (Figure 1). After both sets of expectorate samples were collected, mosquitoes were anaesthetised on ice and bisected following [23]. Head-thorax specimens were stored at -20°C until DNA extraction and subsequent salivary gland sporozoite quantification by qPCR. Only data from sporozoite-infected mosquitoes that survived for the entire experiment were included in all analyses (n=1 *P. berghei* and n=1 *P. chabaudi* uninfected individual mosquitoes were excluded; no uninfected mosquito groups were detected).

**Figure 1.**
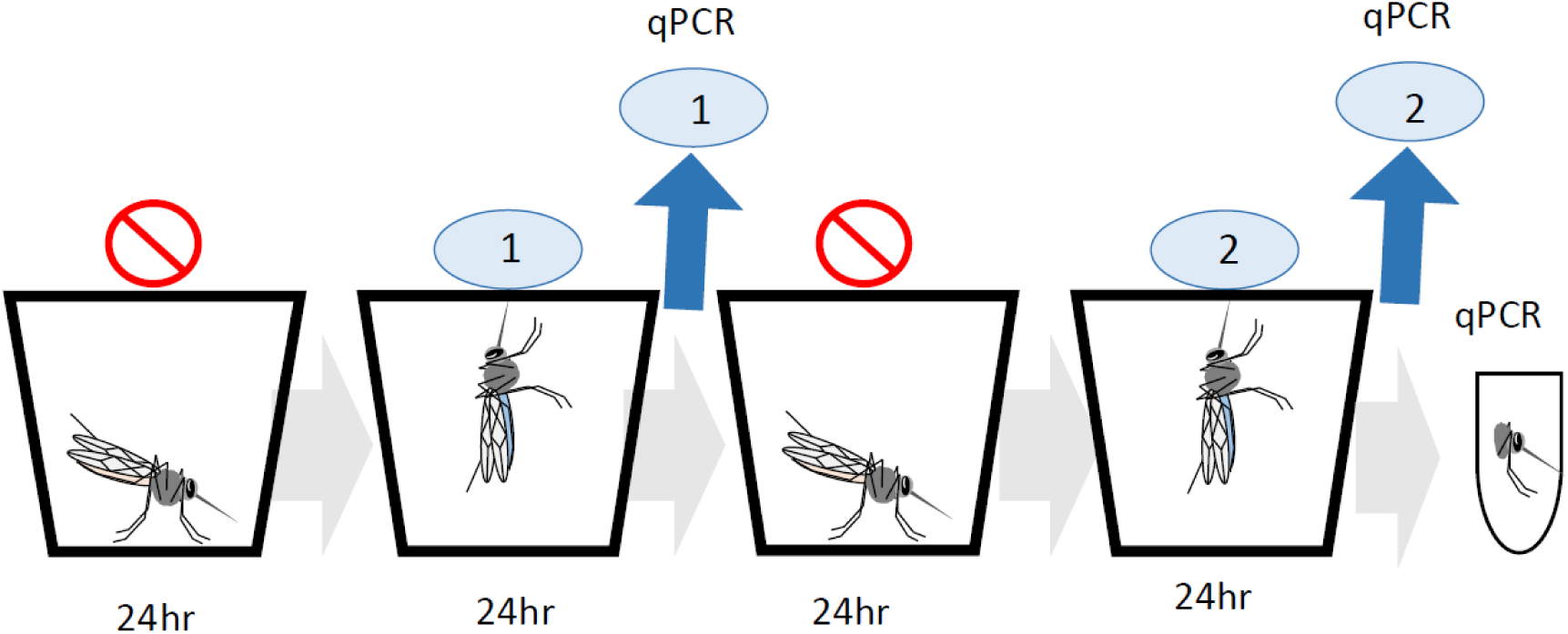
Cotton substrate collection timings from *Plasmodium* infected mosquitoes. Mosquitoes were moved to paper cups and starved for 24 hours, then provided access to a sugar-feeding substrate. After 24 hours, the substrate was collected for sporozoite detection by qPCR. This cycle was repeated twice, such that two substrates were collected per mosquito. The cycle started on day 22 pIBM for *P. berghei* and day 12 pIBM for *P. chabaudi*.

### Statistical analysis

Data analyses were performed using R v. 4.1.3. Linear models were used to determine PCR efficiency, and compare this between species. The absolute limit of detection (LOD) [38], defined as the minimum concentration that can be detected with a sensitivity of 100%, was determined using plateau-linear models fitted to qPCR Ct values and associated genome counts (SSplin, *nlraa* package [39]). These models predict the switching point from a plateau to a linear slope, thus indicating when the qPCR true positivity rate dropped below one. Parasite densities below the LOD were set to zero. To determine the most suitable substrate and sugar concentration, and test for DNA degradation over time, linear models were used to investigate the effect of the variable tested (substrate, sugar, or time), parasites species, DNA concentration (if relevant), and all their interactions on Ct value. DNA yield across *Plasmodium* concentrations was analysed using linear models for *P. chabaudi* and the SSplin function for *P. berghei*, for which this non-linear model fitted better than a linear regression (ΔAICc > 2).

The presence/absence of parasite DNA from expelled sporozoites over time was tested using binomial generalised linear models (glm), including an interaction between *Plasmodium* species and salivary gland burden. Further binomial glms were used to test whether processing a larger proportion of the mosquito expectorate DNA extract (summing parasite densities detected in two qPCR replicates), or collecting expectorates from small groups of *P. berghei-*infected mosquitoes rather than individuals improved detection rates, including species and either replicate or grouping, as well as their interaction into the models. Negative binomial models (glm.nb function, *MASS* package [40]) were used to investigate whether the number of expelled sporozoites on positive substrates was affected by (1) day and salivary gland burden, and how this varied by species; (2) summing parasite densities from two qPCR replicates, by species; (3) grouping *P. berghei*-infected mosquitoes, by day; and (4) whether salivary gland sporozoite burden differed between species.

Models were minimised using likelihood ratio tests, and AICc for non-nested models. All models met model assumptions, confirmed by simulating and plotting residuals using the *DHARMa* package [41]. Confidence intervals were obtained from statistical models or, in the case of confidence intervals for quotients, using Fieller’s method [42].

## Results

### Validation of qPCR for sporozoite detection: True and false positivity

The qPCR assay targeting the 18S rRNA gene has been previously validated for sporozoite detection, achieving a 95% amplification efficiency and a limit of detection of <10 parasites/PCR reaction [36]. We replicate this high qPCR performance using DNA extracted from blood stages of *P. berghei* (0.51 to 6428 genomes/PCR reaction) or *P. chabaudi* (0.24 to 75461 genomes/PCR reaction), achieving an amplification efficiency of 99.5±2.6%, R^2^=0.99, with equal performance between species (log_10_ parasite density by species interaction: *F*_(1,20)_=0.51, *P*=0.482). Although quantification is accurate when low parasite densities are detected, detection rates drop at lower densities. The limit of detection (LOD, the concentration at which the true positivity rate drops below 1) was 4.37 genomes/PCR reaction (Ct 36.7±0.5) for *P. berghei* (Fig 2A) and 0.78 genomes/PCR reaction (Ct 38.4±0.1) for *P. chabaudi* (Fig 2B). At parasite densities below the LOD (i.e. higher Ct values), the rate of false negatives increases and these densities were set to zero. False positives (water samples) were not detected at concentrations above the LOD for either species.

**Figure 2.**
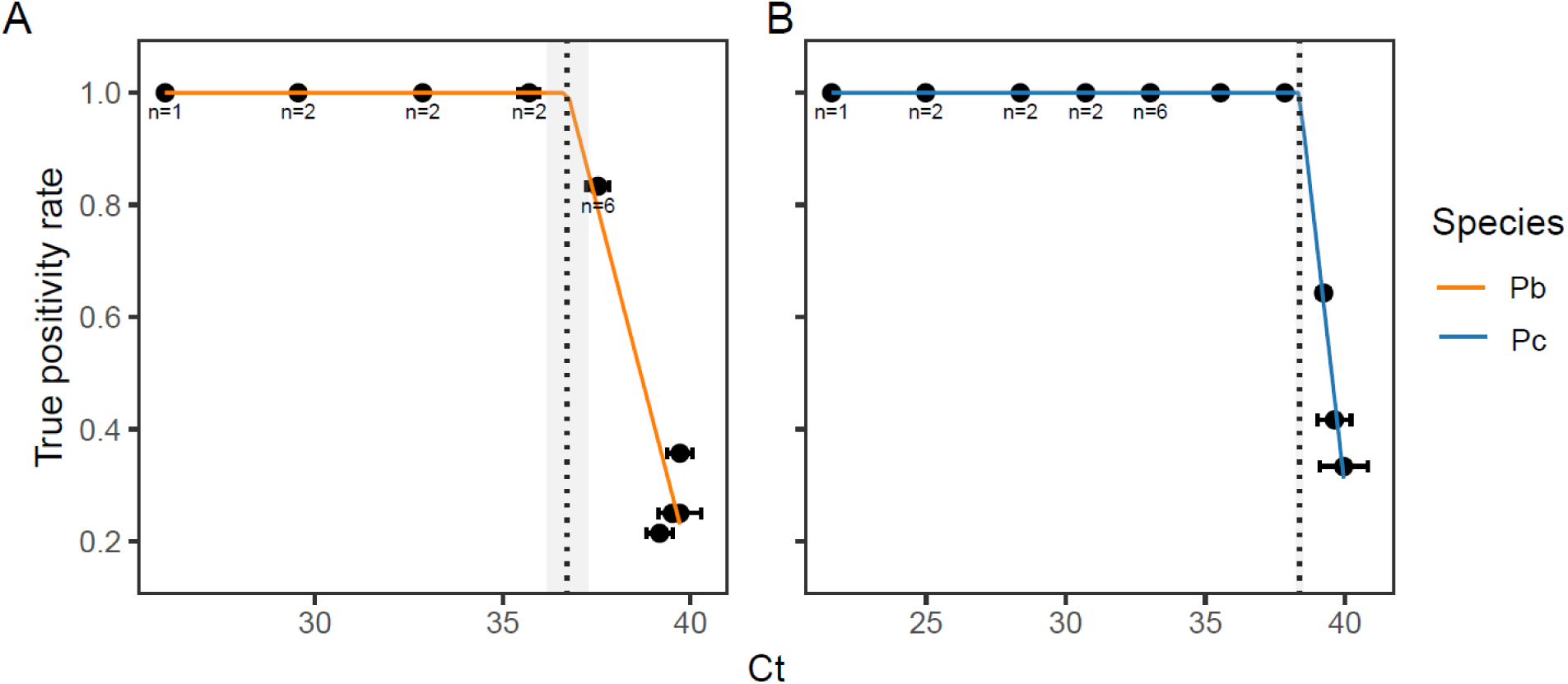
True positivity rates, determined from quantification of a serial dilution of DNA from blood-stage parasites for *P. berghei* (A, orange) and *P. chabaudi* (B, blue). Mean Ct values ±SEM are presented for *P. berghei* (0.001 to 6428 genomes/PCR reaction) and *P. chabaudi* (0.045 to 75461 genomes/PCR), tested in n=12 replicates unless stated otherwise in the graph. The limit of detection (LOD, the concentration at which the true positivity rate drops below 1) ±SEM, predicted using a plateau-linear function, is 4.37 (Ct 36.7±0.5) or 0.78 (Ct 38.4±0.1) genomes/PCR reaction for *P. berghei* and *P. chabaudi*, respectively (dotted lines ± shading).

### Optimising the assay

To determine the most suitable feeding substrate for the assay, three different substrates, soaked in 8% fructose and supplemented with 5µL reference DNA from *P. berghei* or *P. chabaudi* infected mosquitoes, were tested: filter paper, cotton wool and cotton pads. DNA yield varied by substrate type and parasite species (substrate by species interaction: *F*_(2,12)_=4.42, *P*=0.036). Specifically, the extraction efficiency for cotton wool substrates, was 21% and 25% for *P. chabaudi* and *P. berghei* cotton wool substrates, respectively. Cotton pads resulted in DNA yields similar to cotton wool, and filter paper resulted in the lowest DNA yield relative to cotton wool, and more so for *P. chabaudi* (933-fold lower; 95% CI: 186, 4683) than *P. berghei* (276-fold lower; 95% CI: 45, 1675) (Fig 3). Based on DNA yield, and ease of use, cotton wool was selected as the feeding substrate for the remainder of this study.

**Figure 3.**
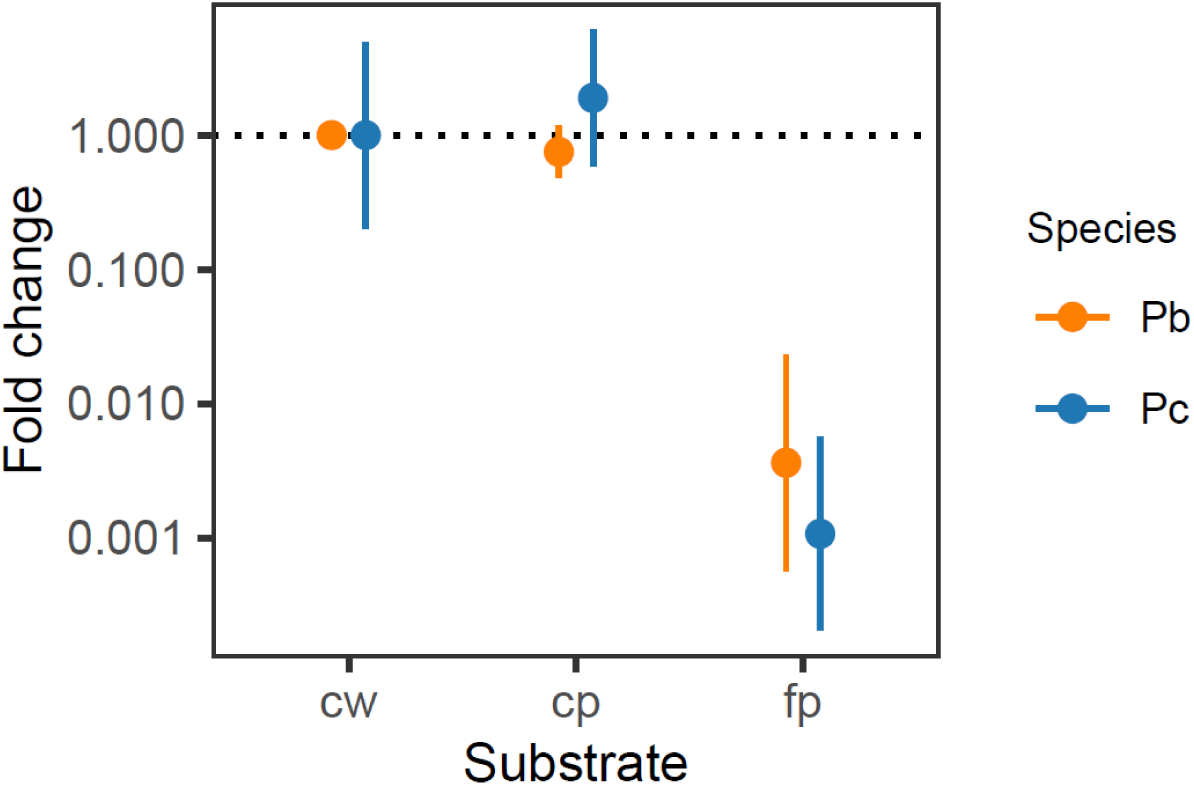
Relative DNA yield from substrates soaked in 8% fructose and supplemented with reference DNA extracted from *P. berghei* (orange) or *P. chabaudi* (blue) infected mosquitoes. DNA yield is presented as mean fold difference ± SEM (n=3/substrates/species) relative to the mean DNA yield for cotton wool for each species and displayed on a log_10_ scale to clearly visualise both increased and decreased DNA yield. Cotton wool (cw); cotton pads (cp), filter paper (fp).

Assay performance for cotton wool was tested by supplementing cotton wool substrates with 5µL of a serial dilution of reference DNA for *P. berghei* (17-3461 genomes/PCR) or *P. chabaudi* (3-5949 genomes/PCR) (Fig 4). Non-linearity for *P. berghei* samples with Ct > 36.5±0.4 shows that quantification became inaccurate at parasite densities below 50 genomes/PCR. This switching point is referred to as the limit of quantification (LOQ), and occurs at a similar Ct value as for *P. berghei* blood samples (36.7±0.5; dotted line, Fig 4A). For *P. chabaudi*, the linear dynamic range covered all tested parasite densities, suggesting that we can confidently detect and quantify *P. chabaudi* genomes from cotton wool substrates up until the LOD as determined by the *P. chabaudi* blood samples above (Ct 38.4±0.1; Fig 4B). The slopes in Fig 4 were steeper than expected, indicating PCR efficiencies of 70.4±8.9% for *P. berghei* and 78.2±4.5% for *P. chabaudi*, which could be explained by covering a wider range of DNA concentrations: DNA quantities were underestimated for low density samples, with Ct values at/above the LOD.To maximise chances of mosquitoes feeding and expelling sporozoites, access to substrates lasted for 24 hrs. Because the conditions in which mosquitoes are kept may not be optimal to preserve DNA, we investigated DNA degradation over 24hrs. Specifically, we compared DNA yield from cotton wool supplemented with DNA at the time of sugar soaking (time point 0) or supplemented at the time of collection 24 hrs later (time point 24). As expected, lower DNA concentrations result in a lower DNA yield (DNA: *F*_(1,33)_=2036.6, *P*<0.001). While the absolute number of genomes varies between species, reflecting the higher parasites densities in the *P. chabaudi* compared to the *P. berghei* reference sample (species: *F*_(1,33)_=310.9, *P*<0.001), quantification is equally efficient in both species (DNA by species interaction: *F*_(1,31)_=0.03, *P*=0.87). We did not observe DNA degradation after 24 hrs of storage for either species (time by species interaction: *F*_(1,29)_=0.0004, *P*=0.98; time: *F*_(1,32)_=3.71, *P*=0.063), across all parasite densities (time by DNA by species interaction: *F*_(1,29)_=1.76, *P*=0.20; time by DNA interaction: *F*_(1,31)_=0.11, *P*=0.75) (Fig 5A).

**Figure 4.**
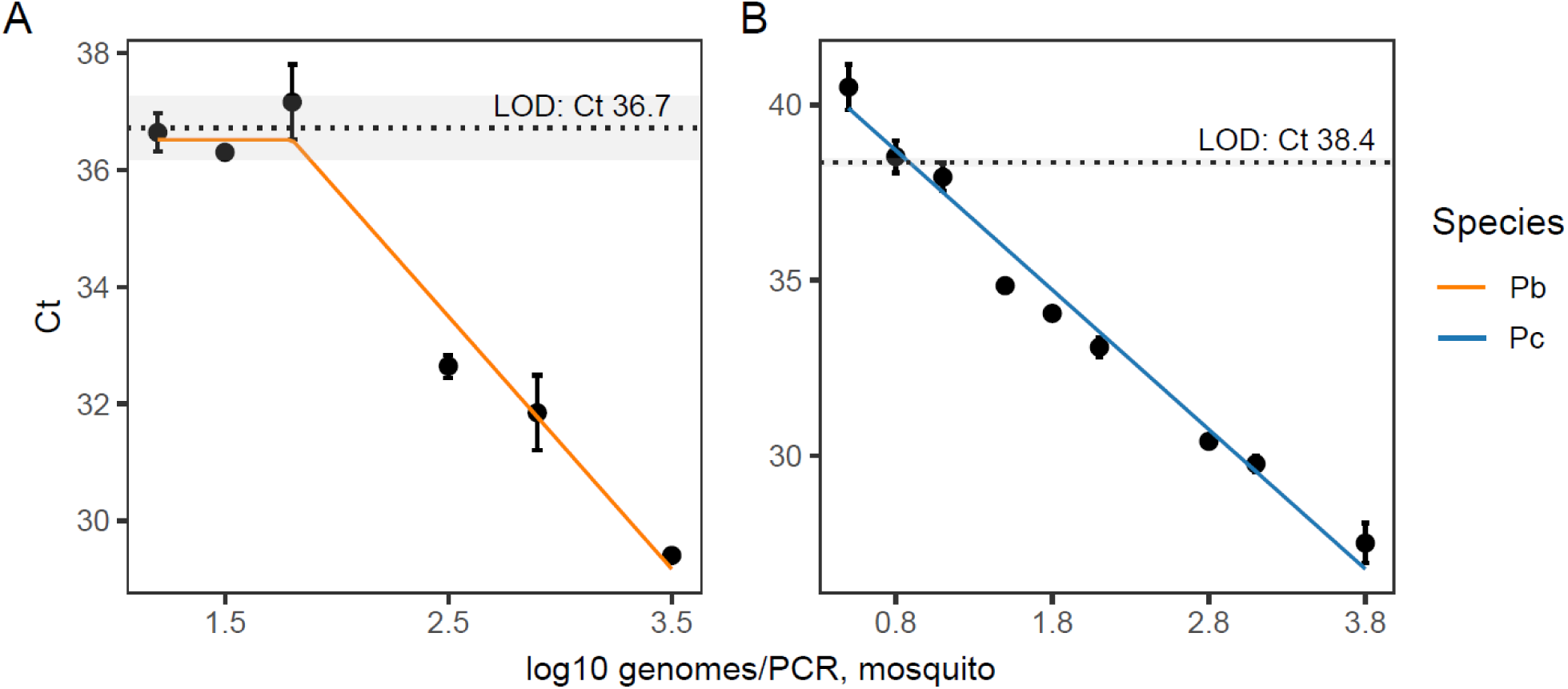
Linearity of quantification for *P. berghei* (orange; A) and *P. chabaudi* (blue; B) reference DNA quantified from cotton wool substrates soaked in 8% fructose. Data points present mean Ct values ± SEM for reference DNA originating from infected mosquitoes (log10 genomes/PCR, mosquito) that ranged from 17-3461 (*P. berghei*) or 3-5949 (*P. chabaudi*) genomes/PCR (n=3 / concentration / species). The limit of quantification (LOQ, Ct at which samples can be successfully detected but quantification becomes inaccurate) is determined as the switch point where the plateau ends. The limit of detection ± SEM (LOD, see Fig 2) and its associated Ct value for blood samples is depicted by the dashed line.

**Figure 5.**
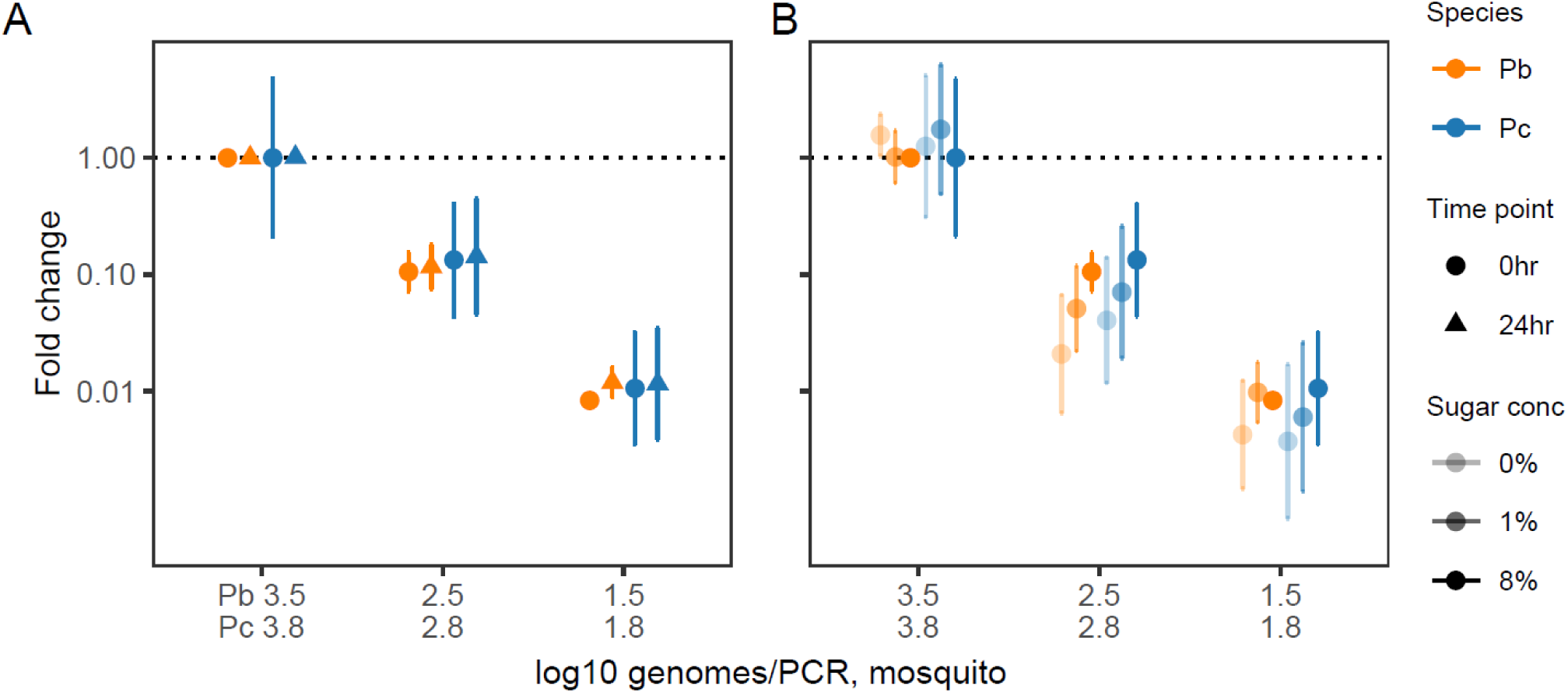
Relative DNA recovery in response to storage time (A) and sugar concentration (B) for reference DNA quantified from cotton wool substrates. Data points present mean fold change ± 95% CI, relative to substrates supplemented with neat reference DNA at time point 0 (A), or soaked in 8% fructose (B), for each species. Dilutions of reference DNA, originating from infected mosquitoes (log10 genomes/PCR, mosq), range from 35-3461 (*P. berghei*, orange) or 59-5949 (*P. chabaudi*, blue) genomes/PCR (n=3 /concentration/group).

Mosquito feeding substrates have a high concentration of sugar (usually fructose or glucose), which may affect extraction efficiency and subsequent amplification of DNA. To test whether the sugar content of the substrates affected DNA yield, we compared DNA recovery from cotton wool substrates soaked in 0, 1 or 8% (w/v) fructose, and supplemented with reference DNA at time point 0. Our analysis confirmed higher parasite densities in the *P. chabaudi*, compared to *P. berghei* reference sample (species: *F*_(1,49)_=80.4, *P*<0.001), that lower DNA concentrations result in lower DNA yields (DNA: *F*_(1,49)_=639.0, *P*<0.001), and that quantification of DNA is equally efficient for both species (DNA by species interaction: *F*_(1,48)_=0.838, *P*=0.36). Sugar concentration impacts DNA yield (sugar: *F*_(2,49)_=8.10, *P*<0.001); lower concentrations reduce the yield compared to 8% fructose by 1.5-fold (95% CI: 1.07, 2.05) for 1% and 2.4-fold (95% CI: 1.72, 3.31) for 0% (Fig 5B), in the same manners across parasite species and densities (sugar by DNA by species interaction: *F*_(2,42)_=0.56, *P*=0.58; sugar by species interaction: *F*_(2,44)_=0.13, *P*=0.88; sugar by DNA interaction: *F*_(2,46)_=1.30, *P*=0.28). Together, these results confirm that collecting mosquito expectorate over a period of 24 hours on substrates soaked in 8% fructose is optimal.

### Testing the assay using mosquito expectorate samples

Following optimisation using reference DNA, we tested the assay’s performance using mosquito expectorate samples. We allowed *Plasmodium-*infected mosquitoes, housed individually (n=12 *P. berghei*, n=9 *P. chabaudi*) or in small groups (n=5 groups of 4 mosquitoes/group for *P. berghei*), to feed for 24 hours on cotton wool substrates soaked in 8% fructose, which were collected twice per (group of) mosquito(es). We compared the prevalence and density of parasite DNA in the feeding substrates between species and by group size.

There was no correlation between the number of sporozoites in the salivary glands of individual mosquitoes and the number of expelled parasites on positive feeding substrates (salivary gland burden: χ^2^_(1)_=0.0098, *P*=0.92) for either species (salivary gland burden by species interaction: χ^2^_(1)_=0.839, *P*=0.36). However, we detected 11-fold (95% CI: 3.61, 34.9) more expelled parasites on positive feeding substrates for *P. berghei* compared to *P. chabaudi* (species: χ^2^_(1)_=10.7, *P*=0.0011; Fig 6A). This likely reflects the 3.4-fold (95% CI: 1.16, 9.56) higher sporozoite burden in the salivary glands for *P. berghei* compared to *P. chabaudi*-infected mosquitoes (species: χ^2^_(1)_=4.50, *P*=0.034; Fig 6B). The number of sporozoites expelled was 2.9-fold (95% CI: 1.32, 6.30) higher on the first vs. second substrate collection day (day: χ^2^_(1)_=4.82, *P*=0.028). This may be due to a higher representation of *P. berghei*-infected mosquitoes, with their higher sporozoite burdens in the glands and the expectorate, in positive substrates of the first (3/4: 75%) vs second collection day (4/6: 67%) (Table 1).

**Figure 6.**
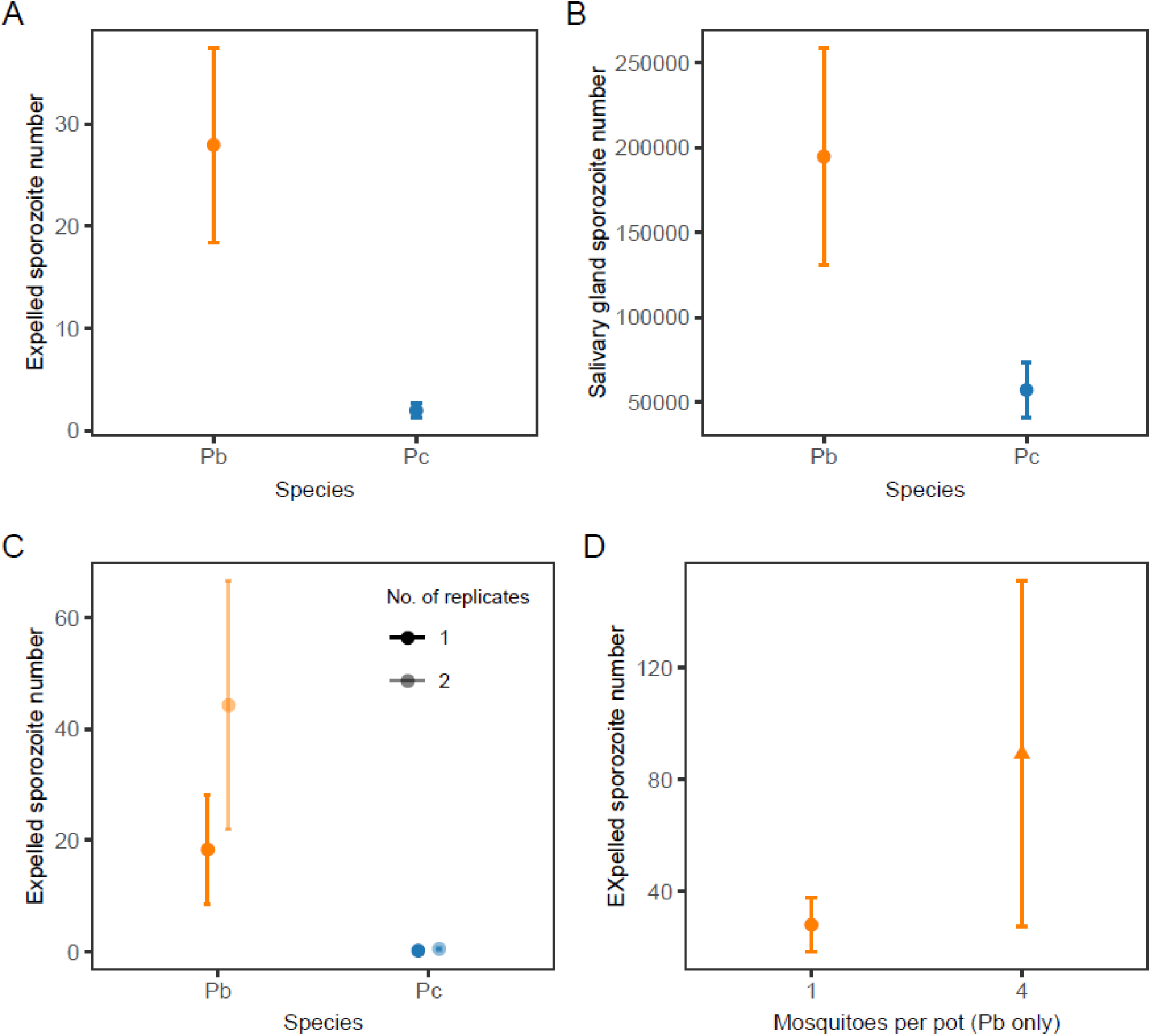
Mean sporozoite densities ± SEM for sporozoite-positive expectorates (A,C,D) and salivary glands (B) for mosquitoes infected with *P. berghei* (orange) and *P. chabaudi* (blue). Data are presented for single (dark colours) or double (C, light colour) qPCR replicates, and for individual mosquitoes (circles) or groups of 4 mosquitoes (triangles, D).

**Table 1.** Parasite detection rates from sugar feeding substrates, fed on by individual or groups of mosquitoes.

|  | *P. berghei* (days 23, 25 pIBM) |  |  | *P. chabaudi* (days 13, 15 pIBM) |  |  |
| --- | --- | --- | --- | --- | --- | --- |
|  | Det. Rate <sup>a</sup> | Ct <sup>b</sup> | Sporozoites<sup>b</sup> | Det. rate <sup>a</sup> | Ct <sup>b</sup> | Sporozoites<sup>b</sup> |
| **Individuals** |  |  |  |  |  |  |
| 1<sup>st</sup> Substrate | 0.25 (3/12) | 33.6±0.6 | 47.4±16.5 | 0.11 (1/9) | 37.9 | 1.5 |
| 2<sup>nd</sup> Substrate | 0.33 (4/12) | 35.2±0.5 | 13.3±4.0 | 0.22 (2/9) | 37.5±0.8 | 2.2±1.1 |
| **Groups** |  |  |  |  |  |  |
| 1<sup>st</sup> Substrate | 0.40 (2/5) | 33.0±2.25 | 172±160 | n.d. | n.d. | n.d. |
| 2<sup>nd</sup> Substrate | 0.60 (3/5) | 34.6±1.16 | 34.4±21.8 | n.d. | n.d. | n.d. |

<sup>a</sup> proportion of positive substrates (number positive/total tested). <sup>b</sup> mean Ct value or number of sporozoites with SEM for positive substrates. n.d : not done

While 50% (6/12) of *P. berghei*- and 33% (3/9) of *P. chabaudi*-infected mosquitoes (confirmed to be positive for salivary gland sporozoites via qPCR) generated at least one positive substrate, the overall proportion of positive feeding substrates from individual mosquitoes was low at 29% (7/24) for *P. berghei* and 17% (3/18) for *P. chabaudi*. Only one mosquito, infected with *P. berghei*, returned a positive substrate on both collection days, and the number of positive substrates was similar across days for both species (day: χ^2^_(1)_=0.63, *P*=0.43) (Table 1). As expected, the probability of detecting parasites on feeding substrates increased with increasing salivary gland burden (χ^2^_(1)_=5.92, *P*=0.015). This was regardless of species, suggesting that parasites from *P. berghei* and *P. chabaudi*, after adjusting for parasite density, are equally well detected in feeding substrates (salivary gland burden by species interaction: χ^2^_(1)_=0.30, *P*=0.58; species: χ^2^_(1)_=0.0044, *P*=0.95).

To investigate whether detection rates of parasites on feeding substrates could be improved, we doubled the proportion of the DNA extract quantified by qPCR from 0.44 to 0.88, by running a replicate qPCR reaction. Doubling the volume of sample tested did increase the density of parasites detected across both replicates in positive cotton substrates by 2-fold (95% CI: 1.03, 5.13) (replicate: χ^2^_(1)_=3.95, *P*=0.047, Fig 6C), regardless of species (species by replicate interaction: χ^2^_(1)_=0.058, *P*=0.809). However, for each species, the proportion of feeding substrates from which parasite DNA was detected was identical (*P. berghei* 7/24, *P. chabaudi* 3/18). Subsequently, we tested if housing mosquitoes in small groups could increase the amount of DNA per feeding substrate (thus improving detection rates), whilst preserving the possibility to obtain data from replicate groups and track results over time. We used groups of 4 *P. berghei*- infected mosquitoes, because the higher sporozoite density in the *P. berghei* expectorates increases the likelihood that sporozoites in the expectorates of multiple mosquitoes will be detectable. As expected, grouping increased the number of expelled sporozoites by 3-fold (95% CI: 1.10, 8.52) (grouping: χ^2^_(1)_=4.00, *P*=0.045; Fig 6D). Less sporozoites were expelled on the second collection day, regardless of group size (day by grouping interaction: χ^2^_(1)_=0.11, *P*=0.74, day: χ^2^_(1)_=6.14, *P*=0.013). However, we did not detect an increase in the rate of detection of *P. berghei* for grouped mosquitoes (grouping: χ^2^_(1)_=1.31, *P*=0.25): 50% (5/10) of cotton substrates obtained from groups returned positive expectorate samples, compared to 29.2% (7/24) from individually-housed mosquitoes (Table 1).

## Discussion

We tested a non-destructive assay for detecting and quantifying sporozoites from mosquito expectorate. Like previous studies [24,26–28], we demonstrate that DNA from *Plasmodium* sporozoites can be detected from feeding substrates. Moreover, we demonstrate our assay detects rodent *Plasmodium* DNA across a range of concentrations, with no evidence of DNA degradation over the 24hrs of sample collection, and DNA yield was optimal on substrates soaked in 8% fructose, which is the concentration commonly used to maintain lab mosquitoes. However, when testing samples expelled by individual mosquitoes (as opposed to reference DNA) the proportion of positive substrates was low.

Our study differs from previous studies in that we investigated individual mosquitoes infected with two commonly used rodent laboratory models, *P. berghei* and *P. chabaudi*. The assay performed similarly for both species and expectorates from mosquitoes infected with *P. berghei* had on average 11-fold more sporozoites than *P. chabaudi-*infected mosquitoes. This is not unexpected considering that we, and others, observe *P. berghei* generally reaches higher oocyst and sporozoite densities [31,43,44], suggesting that there may be malaria species-specific differences in sporozoite inoculum size. Indeed, sporozoite inocula from *P. berghei*-infected mosquitoes are higher than mosquitoes infected with *P. yoelii* [21,45], and more sporozoites are needed to successfully initiate infections in vertebrate hosts for *P. berghei* compared to *P. yoelii,* suggesting that per-sporozoite infectivity is lower for *P. berghei* [46,47]. Although we did not detect a correlation between the number of salivary gland and expelled sporozoites, similar to some studies [20,48–50] but not others [13], we do confirm previous reports [24,25] of higher salivary gland sporozoites burdens correlating with higher rates of detection from substrates. Therefore, like previous studies using *P. falciparum* [19,24,25], our data are consistent with the hypothesis that mosquitoes withhigher salivary gland burdens are more likely to expel sporozoites and transmit malaria.

Low detection rates of expelled sporozoites could be due to technical limitations of the assay. The qPCR limit of detection of approximately four and one genome(s) per PCR for *P. berghei* and *P. chabaudi* respectively, is equivalent to 68 *P. berghei* or 14 *P. chabaudi* genomes (i.e. sporozoites) per substrate when taking into account sample processing and DNA recovery. Additionally, proteins present in mosquito saliva can interact with sporozoites [51], potentially reducing the stability of expelled sporozoites which may raise the detection threshold slightly higher. Therefore, low densities of expelled sporozoites may have gone undetected. However, as detection rates did not improve by running multiple technical replicates, nor for expectorates from groups of four *P. berghei*-infected mosquitoes, it is more likely that not all substrates contain sporozoites. Similar low prevalence of positive feeding substrates were shown in [28] where 31% (day 21 pIBM) and 55% (day 23 pIBM) of cotton wool DNA extracts were positive for groups of three *P. berghei-*infected mosquitoes, in comparison to 40% (day 23 pIBM) and 60% (day 25 pIBM) for our groups of four mosquitoes. Depending on the mosquito species used, 8-52% of feeding substrates contained DNA of the human malaria parasite *P. falciparum* [24]. Our values of 29% and 17% of total positive substrates collected for individually-housed *P. berghei* and *P. chabaudi*-infected mosquitoes respectively sit within this range. In addition, the proportion of individually-housed *An. stephensi* mosquitoes generating at least one positive substrate over 24 hr (33%, *P. chabaudi* and 50%, *P. berghei*) is within the range observed for *P. falciparum* (35%, FTA cards [26]; 61%, artificial skin [13]). Higher proportions of mosquitoes with at least on *P. falciparum* positive cotton substrate (93%) were reported in [24], where up to ten substrates per mosquito were collected, thus increasing the chance of at least one substrate being positive. Together these data indicate that the low detection rate of parasites on feeding substrates may be common.

Low detection rates on feeding substrates may instead be due to mosquito feeding behaviour, sporozoite biology or a combination of both. While mosquitoes do not expel sporozoites every day [24,25,28], female mosquitoes are likely to sugar-feed daily especially if they do not have access to a blood meal [52]. As mosquitoes in our study were starved for 24 hours in between access to substrates to increase feeding rate, a lack of sugar-feeding is unlikely to explain the absence of sporozoites on the feeding substrates. It is generally assumed that mosquitoes salivate in a similar way during sugar- and blood-feeding [26,30], although different enzymes are released from different lobes during the two types of feeding, and thus sporozoite expulsion may vary too. Furthermore, when mosquitoes with high *P. yoelii* sporozoite loads were blood-fed on multiple days, some did not inject any sporozoites and some only injected a high number of sporozoites on one of the days [21]. This may be due to sporozoites clumping together [53,54], which will increase variation in expulsion probability. We found that *P. berghei* sporozoite expulsion was lower on day 25 than on day 23 pIBM, and others did not detect any *P. berghei* sporozoites from 26 days pIBM onwards [28]. This suggests that sporozoites may degenerate [55] or deplete [56] over time. If so, *Plasmodium* species may vary in sporozoite lifespan in the glands; for example, *P. falciparum* sporozoites have been shown to be expelled for several weeks [24].

How the quantity and quality of sporozoites (both in the salivary glands and expectorate) influences the probability of transmission remains mysterious. While expelled sporozoites in this assay cannot be directly tested for their infectivity to hosts, patterns of sporozoite expulsion over time may provide a better proxy for mosquito infectivity to vertebrate hosts than the presence or density of salivary gland sporozoites at a certain time point. Expelled sporozoites are transcriptionally different to those in the glands [14], and thus may vary in their properties, including infectivity. Therefore, a more appropriate measure of EIP may be the time at which sporozoites are first expelled, rather than when sporozoites appear in the salivary glands. However, our data suggest that to estimate infectivity throughout a mosquito’s lifespan, the frequency of expulsion should be accounted for as well. While our assay can detect expelled sporozoites from sugar feeding substrates for two rodent malaria species, further assay improvements are needed to track EIP over time in individual mosquitoes. Identifying why detection rates are low remains a key challenge for improving the assay. Our qPCR assay has high sensitivity, so increasing DNA recovery is most likely to improve detection rate. For example, a liquid-only feeding system could reduce yield losses and therefore increase assay sensitivity. Additionally, setting up video systems or using food colouring in sugar substrates would provide further information on the frequency of mosquito feeding. Confirming how often mosquitoes feed would allow untouched negative substrates to be excluded, and is also key to resolving the likelihood of sporozoite expulsion over time.

## Conclusions

Rodent malaria species are a valuable laboratory tool for comparison between different *Plasmodium* species and for asking broad questions about *Plasmodium* biology. We show that expelled sporozoites from two different rodent malaria species can be detected from feeding substrates, but further improvement is needed to use this assay for tracking sporozoite expulsion from individual mosquitoes. The low rate of parasite detection in feeding substrates suggests that the appearance and burden of salivary gland sporozoites may not be the most appropriate measure of mosquito infectivity, and that the definition of EIP may require updating. Tracking expelled sporozoites in individual mosquitoes, rather than using salivary gland sporozoite dissections, would be optimal, whilst facilitating studies to identify how environment-parasite-vector interactions influence EIP and infectivity to vertebrate hosts over time.

## Acknowledgements

We thank Aidan O’Donnell, Ronnie Mooney and Jacob Holland for practical help, and Anna Cohuet and Edwige Guissou for discussion and suggestions to improve the assay.

## Author contributions

Conceptualization, PS, SER and CEO; Investigation, CEO; Analysis, CEO, PS; Writing-Original Draft, CEO, PS; Writing-Review and Editing, all authors.

## Availability of data and materials

The datasets supporting the conclusions of this article are available in Edinburgh DataShare repository [http:// ADD LINK UPON ACCEPTANCE]

## Ethics approval and consent to participate

All procedures comply with the UK Home Office regulations (Animals Scientific Procedures Act 1986; SI 2012/3039) and were approved by the ethical review panel at the University of Edinburgh (PPL PP8390310).

## Consent for publication

Not applicable.

